# CLIC4 is a cytokinetic cleavage furrow protein that regulates cortical cytoskeleton stability during cell division

**DOI:** 10.1101/723940

**Authors:** Eric Peterman, Mindaugas Valius, Rytis Prekeris

## Abstract

During mitotic cell division, the actomyosin cytoskeleton undergoes several dynamic changes that play key roles in progression through mitosis. While the regulators of cytokinetic ring formation and contraction are well-established, proteins that regulate cortical stability during anaphase and telophase have been understudied. Here, we describe a role for CLIC4 in regulating actin and actin-regulators at the cortex and cytokinetic cleavage furrow during cytokinesis. We first describe CLIC4 as a new component of the cytokinetic cleavage furrow that is required for successful completion of mitotic cell division. We also demonstrate that CLIC4 regulates the remodeling of sub-plasma membrane actomyosin network within the furrow by recruiting MST4 kinase and regulating ezrin phosphorylation. This work identifies and characterizes new molecular players involved in the transition from the contracting cytokinetic ring to the intercellular bridge during cytokinesis.

## Introduction

Cytokinesis is the last step in mitotic cell division that relies on orchestrated changes in membrane traffic and cytoskeletal rearrangements required to complete the separation of the two daughter cells. Failure in cytokinesis has wide-ranging effects, including apoptosis and ploidy. Understanding these cytoskeletal rearrangements and shape changes is paramount to understanding cell division during development and disease contexts such as cancer. A number of highly controlled shape changes owing to these cytoskeletal rearrangements occur during cell division, from rounding up in metaphase, to furrowing in anaphase, and finally to the formation of the intercellular bridge in telophase. During metaphase, mitotic rounding contributes to creating the geometric space necessary for spindle organization and morphometry ^1^. Some of the proteins that are well-known to regulate these changes in metaphase include non-muscle myosins IIA/B (NMYIIA/B), as well as the members of ezrin-radixin-moesin (ERM) family of proteins. They do so by regulating actin dynamics at the cell cortex, where they assist in maintaining cortical actomyosin network and rigidity of the plasma membrane. Loss of these proteins leads to altered shape changes and oftentimes a failure in cell division ^2–5^.

Following metaphase, cells undergo the process of cytoplasmic separation, also known as cytokinesis. Cytokinesis relies heavily on the formation and contraction of the cytokinetic actomyosin ring, an actin-rich structure that forms at telophase onset. A number of proteins have been shown to be required for cytokinetic ring formation and contraction, including RhoA, anillin, and non-muscle myosin family members ^6–9^. RhoA is a key regulator of this process since its activation drives formation and constriction of the cytokinetic ring through an increase in actin polymerization and myosin activation. Anillin is a scaffolding protein that is contributes to regulating the localization of RhoA, actin and septins to the contractile ring. While furrows can initiate in the absence of anillin, cytokinesis often fails due to the necessity of its scaffolding partners.

An important consideration in telophase during furrow ingression is that the cortexes of the dividing cells remain mostly rigid, while the furrow is constricting to separate the cytoplasm. These local changes in stiffness are regulated by the actomyosin cytoskeleton in conjunction with the ERM proteins, and ultimately aid in furrow ingression. An additional facet of this regulation are the pressure dynamics induced by the rigidity of the cell cortex. It is thought that self-induced ruptures and blebs of the cortex allow for pressure release and for cell division to continue. However, disruption of the actomyosin cortex via manipulation of these proteins can lead to abnormal blebbing events, which can lead to a failure in cell division ^10–12^. While nonmuscle myosins and ERM proteins have been extensively studied and are well-known to regulate this blebbing phenomena, it is still poorly understood how these blebs are regulated during cytokinesis.

Upon completion of furrow ingression, the daughter cells remain connected by a thin and actin rich intracellular bridge (ICB). The conversion of the contracting actomyosin ring to a stable, actin-rich ICB is a key step in mitotic cell division, and failure in this transition often leads to furrow regression and polyploidy. However, the molecular machinery mediating the conversion of formin-based actomyosin ring to stable sub-plasma membrane actin network in ICB remains largely unknown. In our previous analysis of the midbody proteome, we identified a number of proteins that have the potential to regulate actin cytoskeletal dynamics during cell division. One such protein is CLIC4. CLIC4 (chloride intracellular channel 4) is a 28kDa protein that belongs to the family of CLIC proteins. As its name might imply, it was initially described as a chloride channel; however, numerous recent studies have shown that CLIC4 may actually function as a regulator of actin dynamics. These studies implicated CLIC4 in regulating branched actin networks and filopodia formation, although how CLIC4 performs these functions remains to be defined ^13–16^. In this study, we describe a role for CLIC4 in ensuring the fidelity of cell division through the regulation and recruitment of actin-interacting proteins during cell division. We show that CLIC4 is recruited to the cytokinetic furrow by RhoA at the onset of furrow formation. We also demonstrate that CLIC4 binds to MST4, a kinase and known regulator of ERM proteins, and mediates recruitment of MST4 to the ingressing furrow during telophase. The absence of CLIC4 results in decreased recruitment of phospho-ezrin and non-muscle myosin IIA/B to the furrow, leading to increased membrane blebbing and a failure to transition from cytokinetic actomyosin ring to ICB. Collectively, our data demonstrate that CLIC4 is a new component of the contractile cytokinetic ring that contributes to regulating cortical actomyosin dynamics during cell division.

## Materials and Methods

### Cell culture and treatments

HeLa and MDCK cells were kept in 37° humidified incubator at 5% CO2, routinely tested for mycoplasma, and were maintained in DMEM with 5% FBS and 1% penicillin/streptomycin. To create the GFP-CLIC4 cell line, HeLa or MDCK cells were transduced with the lentivirus pLVX:GFP-CLIC4. Populations were selected with puromycin.

To create shCLIC4 stable knockdown cell lines, HeLa cells were transduced with the lentivirus TRCN0000044358. Populations were selected with puromycin, and stable clones were isolated. For cell synchronization, a double thymidine-nocodazole block was used. Cells lysates were harvested when cells entered anaphase. For RhoA inhibition, Rho Inhibitor I (Cytoskeleton) was used at a final concentration of 0.5ug/ml for 6 hours. IAA94 was used at a concentration of 50 μM for 24 hours. The anillin siRNA sequence is as follows: CGAUGCCUCUUUGAAUAAAUU and was used at a final concentration of 250nM.

To knock-out CLIC4, MDCK cells expressing dox-inducible Cas9 were co-transfected with two CLIC4 gRNAs (GCUGUCGAUGCCGCUGAAGUUUUAGAGCUAUGCUGUUUUG and UGACGAAGAGCUCGAUGAGUUUUAGAGCUAUGCUGUUUUG). Cells were then plated at low density and resulting clones were isolated and tested for presence of CLIC4 by western blot.

f

### GFP-tagging endogenous CLIC4

The Horizon Discovery Edit-R system was used to insert GFP into the endogenous CLIC4 locus. Briefly, 5- and 3- homology arms were PCR amplified and a homology-directed repair template with GFP was created using Gibson Assembly. GFP was inserted into the endogenous locus immediately downstream of the ATG start site. HeLa cells expressing Cas9 under a doxycycline-inducible promoter were transfected with the HDR plasmid and with gRNA CAAGUUCUGCACAGGUCUGCGUUUUAGAGCUAUGCUGUUUUG, flow sorted, and CLIC4 was sequenced and probed for via western blot. To confirm the insertion of the GFP tag at the N-terminus of endogenous CLC4 allele, all HeLa lines were genotyped by sequencing.

### Plasmids and antibodies

The following antibodies and reagents were used for immunofluorescence: anti-acetylated tubulin (Sigma, T7451) Hoechst 33342, SiR-actin (Cytoskeleton, Inc), anti-phospho-ERM (CST 3726), anti-MST4 (Proteintech 10847), anti-NMYIIA (Abcam 75590), anti-NMYIIB (CST 3404), anti-anillin (EMD Millipore MABT96), anti-RhoA (Santa Cruz sc-418). AlexaFluor-594- and AlexaFluor-488-conjugated anti-rabbit and anti-mouse secondary antibodies were purchased from Jackson ImmunoResearch Laboratories (West Grove, PA). AlexaFluor-568-phalloidin was purchased from Life Technologies (Carlsbad, CA). The following antibodies were used for western blotting: anti-CLIC4 (Santa Cruz sc-135739), anti-alpha-tubulin (sc-23948), anti-MST4 (Proteintech 10847). The IRDye 680RD Donkey anti-mouse and IRDye 800CW donkey anti-rabbit secondary antibodies used for western blotting were purchased from Li-COR (Lincoln, NE). The following plasmids were used: pCAG:IRES-CLIC4 (gift from Sung Lab, Cornell), pEGFP-N3:CLIC4, RhoA FRET biosensor (gift from Bokoch), pGEX:CLIC4, pCMV-SNAP-CRY2-VHH(GFP) (Addgene #58370), Tom20-CIB-stop (Addgene #117243), Cry2PHR-mCH-RhoA (Addgene #42959), and pGEX6P1-GFP-Nanobody (Addgene #61838. Site directed mutagenesis was used to create pEGFP-N3:CLIC4C35A.

### Microscopy

Routine imaging was performed on an inverted Axiovert 200M microscope (Zeiss) with a 63× oil immersion lens and QE charge-coupled device camera (Sensicam). Z-stack images were taken at a step size of 500-1000 nm. Image processing was performed using 3D rendering and exploration software Slidebook 5.0 (Intelligent Imaging Innovations) or in ImageJ. Images were not deconvolved unless otherwise stated in the figure legend. Time lapse imaging was performed using a Nikon A1R, equipped with a humidified chamber and temperature-controlled stage. Time between images is displayed in the figures of each experiment.

### Optogenetics for Mitotrap and FRET analysis

For the RhoA activation and GFP-CLIC4 recruitment assay, cells expressing GFP-tagged endogenous CLIC4 were transfected with Cry2PHR-mCH-RhoA (Addgene #42959). Cells were kept in the dark and incubated for 24 hours. To induce clustering and activation of RhoA, cells were exposed to 488 nm light for 5 seconds with the laser power varied between 1 and 10%.

For the Mitotrap experiments, cells expressing endogenous GFP-CLIC4 were transfected with Cry2-VHH Addgene #58370) and Tom20-CIB-stop (Addgene #117243). Cells were kept in the dark and incubated for 24-48 hours. To induce mitochondrial anchoring of GFP-CLIC4, cells were exposed to 488nm light for 5 seconds with the laser power varied between 1 and 10%. For fixation, cells were fixed ~20 minutes following 488nm light pulse. For the Mitotrap time-lapse assays, mitochondrial anchoring was ensured for the entirety of the experiment as cells were pulsed and imaged every 5 minutes.

### Identification of CLIC4-interacting proteins

To identify CLIC4-interacting proteins, cells expressing GFP alone or pLVX:GFP-CLIC4 were lysed and proteins were immunoprecipitated using a GST-tagged GFP-nanobody (Addgene #61838) that was covalently linked to Affigel 10/15 resin (BioRad). First, lysates were precleared with GST linked to Affigel, then incubated with GFP-nanobody-containing Affigel. Affigel beads were then washed and resuspended in 500 μL of 50 mM ABC to remove residual DTT and IAA. After centrifugation and supernatant removal, ProteaseMax (0.02%) and MS-grade trypsin was added (1:50) to the particle pellet. The beads were then washed twice in 100 ul of 80 % CAN with 1 % FA and the supernatants were pooled and analyzed by LC-MS. Scaffold (version 4.8, Proteome Software, Portland, OR, USA) was used to validate MS/MS based peptide and protein identifications. Peptide identifications were accepted if they could be established at greater than 95.0% probability as specified by the Peptide Prophet algorithm. Protein identifications were accepted if they could be established at greater than 99.0% probability and contained at least two identified unique peptides. The criteria to determine CLIC4-interacting proteins were to use a 1.5-fold enrichment cutoff of spectral counts in GFP-CLIC4 over empty GFP (Supplemental Table 1).

For GST pulldown assays, GST-CLIC4 was purified from BL21-Codon PLUS E.Coli as previously described ^17^. HeLa cells were lysed in buffer containing 20mM HEPES, 150mM NaCl, protease and phosphatase inhibitors and 0.1% Triton X-100. 1mg of HeLa cell lysate was mixed with 10μg of either GST or GST-CLIC4 and incubated for 30 minutes at room temperature. Glutathione beads were added and incubated for an additional 60 minutes. Beads were washed, protein was eluted with 1% SDS and separated by SDS-PAGE, followed by western blot analysis.

### Image quantification and Statistical analysis

Unless described otherwise, statistics were performed using a Student’s t-test. For multinucleation assays, at least 5 random fields were selected and imaged. For measurements containing cells in anaphase at least 15-30 cells were randomly chosen and imaged. For measuring cell division times, metaphase onset was determined as cell rounding and conclusion of telophase/division was determined as cell flattening and no apparent intercellular bridge. Fluorescent intensities were measured using 3i software or ImageJ and displayed either as raw fluorescence or as fluorescence ratio (poles versus furrow). In all cases data were collected from at least from three independent experiments.

## Results

### CLIC4 is enriched at the cytokinetic furrow

We have recently completed a proteomic analysis of post-mitotic midbodies purified from HeLa cell media ^18^. Post-mitotic midbodies are released into the media as a result of symmetric abscission, and therefore contain the membranous envelope derived from the plasma membrane of the intracellular bridge connecting two daughter cells during telophase (Figure 1A). Consequently, post-mitotic midbodies are also expected to contain proteins that regulate cytokinetic actomyosin ring contraction during early telophase and disassembly during abscission. Consistent with this idea, numerous cytokinesis regulators, such as RhoA, MKLP1, Plk1, anillin, Rab11 and Rab35 were all identified in midbody proteome ^18–20^. Among these known cytokinesis regulators, CLIC4 was also identified as a potential midbody-localizing protein ^18,21^. Since CLIC4 was previously implicated in regulating actin cytoskeleton dynamics during cell migration, we hypothesized that CLIC4 may also be involved in regulating actin networks during cytokinesis. To test this hypothesis, we first analyzed the localization of CLIC4 during cytokinetic ring formation and contraction. To that end we used a fluorescent protein-tagged version of CLIC4 (N-terminus). As previously reported ^13^, GFP-CLIC4 predominantly localizes at the plasma membrane as well as actin filaments at both the plasma membrane cortex and in filopodia during interphase (Supplemental Figure 1A). Consistent with the involvement of CLIC4 in regulating cytokinesis, CLIC4 is enriched at the cleavage furrow and the contractile ring in early and late anaphase (Figure 1B, D and F, Supplemental Figure 1, Supplemental Movie 1, Supplemental Movie 2). To ensure that localization of GFP-CLIC4 is not a result of overexpression, we next used CRISPR/Cas9 to tag endogenous CLIC4 with GFP on the N-terminus (Supplemental Figure 1B). As shown in Figure 1E-F, endogenously tagged CLIC4 was also targeted to the cleavage furrow during anaphase where it colocalized with anillin, one of the well-established contractile ring proteins. Interestingly, when cells expressing endogenous GFP-CLIC4 were treated with siRNA targeting anillin, we saw a slight but significant decrease in GFP-CLIC4 levels at the contractile ring, suggesting that anillin may contribute to regulating CLIC4 localization during anaphase (Supplemental Figure 1C).

**Figure 1.**
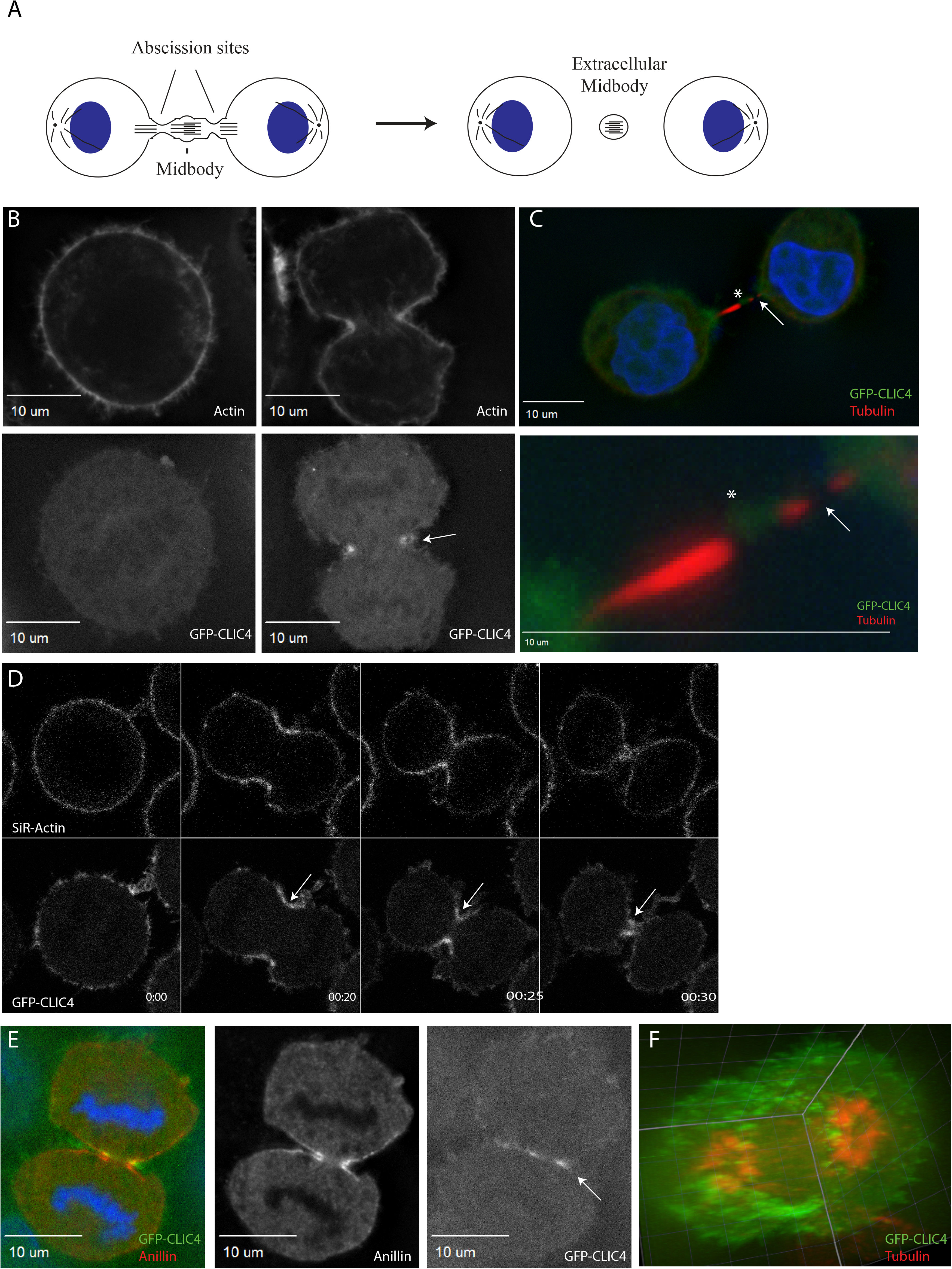
Localization of CLIC4 throughout the cell cycle. (A) Schematic representation of abscission that generates extracellular post-mitotic midbody. (B) HeLa cells expressing exogenous GFP-CLIC4 were fixed and stained with phalloidin-Alexa568 (left and center images) or anti-acetylated tubulin antibodies (right image). Arrow in center image points to ingressing cleavage furrow. Arrow in center image points to abscssion site. Asterisk marks the midbody. (C) Hela cells expressing exogenous GFP-CLIC4 were analyzed by time lapse microscopy. Images shown are stills representing different sequential time points during anaphase. Arrow marks cleavage furrow. (D) HeLa cells expressing endogenously tagged GFP-CLIC4 were fixed and stained with anti-anillin antibody. Arrow points to the cleavage furrow. (F) 3D volume rendition of GFP-CLIC4 expressing Hela cell in anaphase. Red marks central spindle microtubules labeled with anti-acetylated tubulin antibodies.

During telophase and upon completion of cytokinetic ring contraction, GFP-CLIC4 remained associated with the intercellular bridge and the midbody (Figure 1D), explaining its identification in midbody proteome ^18^ and consistent with a previous report suggesting CLIC4 is a midbody protein ^21^. During abscission, GFP-CLIC4 is removed from the abscission site (Figure 1C, lower right panels). This observation is consistent with previous reports that actin filaments needs to be removed from the abscission site to allow for recruitment of the ESCRT-III complex and final separation of daughter cells ^22,23^. Collectively, we propose that CLIC4 is a newly described component of the cytokinetic furrow that may regulate the actomyosin ring contraction, as well as formation and stabilization of the intracellular bridge (ICB).

### RhoA is necessary and sufficient to localize CLIC4 at the cytokinetic cleavage furrow

Since previous studies have reported that RhoA regulates CLIC4 localization in interphase cells ^13^, we hypothesized that activated RhoA at the cytokinetic furrow may be responsible for targeting of CLIC4 during cytokinesis. To test this hypothesis, we first confirmed that CLIC4 indeed colocalizes with active RhoA during anaphase. Using a TCA fixation method that is known to preserve RhoA localization during cell division and anti-RhoA antibody, we observed endogenous GFP-CLIC4 colocalizing with RhoA (Figure 2A). Next, we tested if RhoA activation is sufficient to drive CLIC4 relocalization. To do so, we used our endogenous GFP-tagged CLIC4 cell line in tandem with an optogenetic RhoA clustering assay ^24^. To that end, we used HeLa cells expressing Cry2mCH-RhoA. The exposure of cells to 488nm wavelength light leads to clustering and activation of RhoA within the cytoplasm ^25^(see Methods for detailed description). As shown in Supplemental Figure 1D, endogenous GFP-CLIC4 accumulated in light-induced activated RhoA complexes, suggesting that RhoA activation is sufficient to recruit CLIC4.

**Figure 2.**
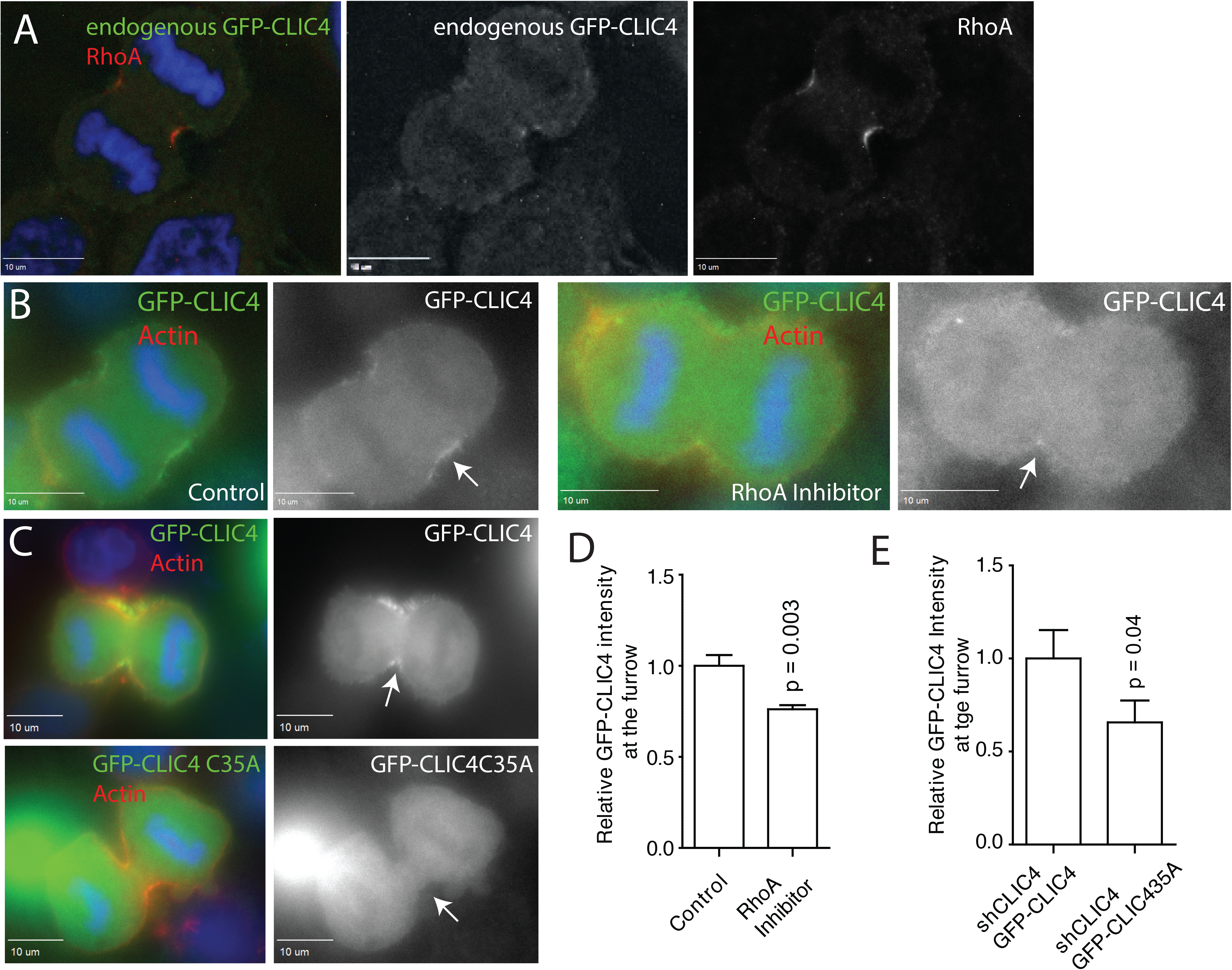
RhoA is necessary and sufficient for CLIC4 localization. (A) HeLa cells expressing endogenous GFP-CLIC4 were TCA-fixed and stained with RhoA antibodies. Image was deconvolved. (B) Hela cells expressing endogenously labeled GFP-CLIC4 were imaged in anaphase in the presence or absence of RhoA inhibitor. Arrows point to cleavage furrows. (C) shCLIC4 HeLa cells were transfected with either wild-type GFP-CLIC4 or GFP-CLIC4 C35A. Cells were then fixed and stained with phalloindin-Alexa568. Arrows point to cleavage furrow. (D-E) Quantification of GFP-CLIC4 (D and E) or GFP-CLIC4 35A (E) fluorescence in cleavage furrow. Data shown are the means and standard deviations of 15-30 anaphase cells randomly picked during three independent experiments.

We next tested if RhoA is necessary for the recruitment of CLIC4 to the cytokinetic ring. First, we used a low dose (0.5μg/ml for 4 hours) of a Rho inhibitor that still allowed furrow formation, although at reduced rates and efficiency. Consistent with the involvement of RhoA in mediating CLIC4 recruitment, RhoA inhibitor decreased the levels of endogenous GFP-CLIC4 at the furrow (Figure 2B and D). Second, we examined the localization of GFP-CLIC4-C35A, a mutant which was previously shown to lack the ability to translocate to sites of RhoA activation ^13^. GFP-CLIC4 or GFP-CLIC4-C35A were expressed in CLIC4-knockdown HeLa cells (shCLIC4), and fluorescent intensity was measured and compared at the contractile ring as well as at the poles of the dividing cell. As shown in Figure 2E, GFP-CLIC4-C35A was not enriched at the contractile ring as compared to GFP-CLIC4 (also see Figure 2C), consistent with RhoA involvement in recruiting CLIC4 to the furrow during anaphase. Together, these data suggest that RhoA is responsible for CLIC4 recruitment to the ingression furrow during anaphase.

### CLIC4 is required for actin-dependent stabilization of the intracellular bridge

Our data demonstrate that CLIC4 is recruited to the plasma membrane at the cytokinetic furrow during telophase and remains associated with intracellular bridge during abscission. However, the function of CLIC4 remains unclear. We next asked if CLIC4 is required for any aspect of cytokinesis. To determine that, we created a HeLa cell line stably expressing CLIC4 shRNA (shCLIC4) (Supplemental Figure 2A-B). As shown in Figure 3A, CLIC4 knockdown resulted in an increase in multi-nucleation in HeLa cells, indicating defects in cytokinesis. In contrast, knock-down of another closely related CLIC family member, CLIC3, did not induce a multi-nucleation phenotype (Figure 3A). Finally, this increase in multi-nucleation could be rescued by re-introduction of GFP-CLIC4, but not GFP-CLIC4-C35A (Figure 3A). It is also worth noting that, despite extensive effort, we were unable to create a viable CLIC4 knock-out HeLa cell line using CRISPR/Cas9, suggesting that complete deletion of CLIC4 may block cell division in HeLa cells. In contrast, our shCLIC4 cell lines still expresses ~5% of CLIC4, allowing cells to eventually complete cytokinesis.

**Figure 3.**
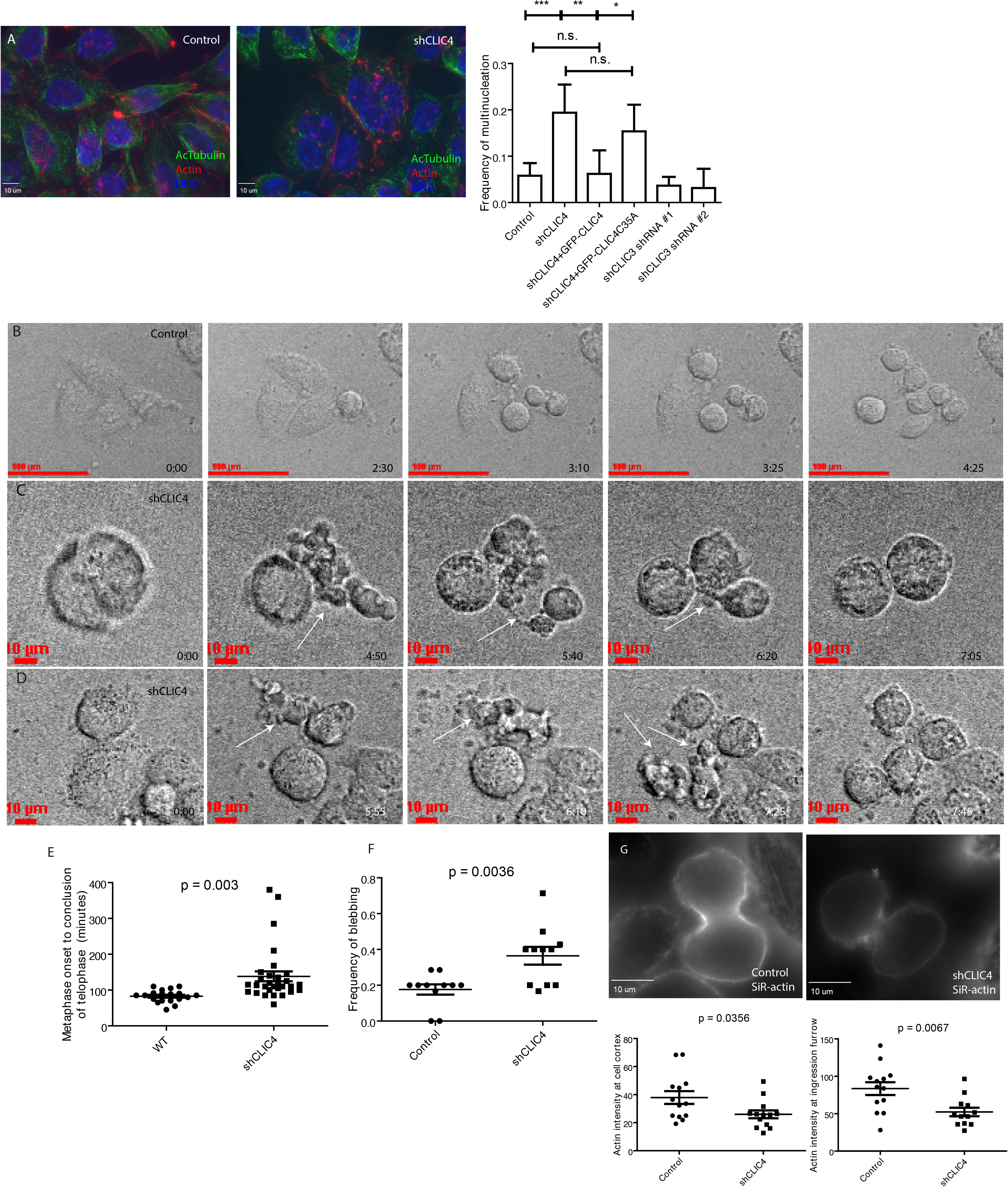
Depletion of CLIC4 leads to mitotic defects. (A) HeLa cells were fixed and co-stained with phalloidin-Alexa568 and anti-acetylated tubulin antibodies. The number of multi-nucleated cells were then counted. Graph on the right shows quantification and statistical analysis of multi-nucleation induced by shRNA-dependent CLIC4 depletion. Data shown are the means and standard deviations of at least 75 cells counted in randomly chosen fields from three different experiments. (B-D) Control (B) or shCLIC4 (C and D) HeLa cells were analyzed by time lapse microscopy. Images shown are the stills from different time points of cells undergoing mitotic cell division. Arrows point to blebs during anaphase (E) Quantification of time required for cells to go from metaphase to abscission. Data shown are means and standard deviations derived from randomly picked cells in two different experiments. (F) Quantification of blebbing frequency in control or shCLIC4 Hela cells. Data shown are means and standard deviations derived from randomly picked cells in three different experiments. (G) Control or shCLIC4 cells were incubated with SiR-actin and live imaged. Quantification of actin intensity at the cell cortex and furrows are shown. Data shown are individual cell intensities from randomly picked cells in three different experiments.

Since CLIC4 was initially described as a putative chloride ion channel, we next tested the effect of IAA94, a chloride channel inhibitor, on cell division. Treatment of cells with this inhibitor did not induce multi-nucleation (Supplemental Figure 2C), nor did it affect CLIC4 recruitment to the furrow (Supplemental Figure 2D), suggesting that, at least in the context of cell division, CLIC4 does not function as chloride channel.

To determine what part of cytokinesis depends on CLIC4, we next assessed cell division using time-lapse microscopy. As shown in Figure 3E, CLIC4 depletion led to an increase in the amount of time it took cells to complete mitosis (also see Supplemental Movies 4 and 5). While all shCLIC4 cells formed an ingression furrow, about 20% of furrows regressed at late anaphase resulting in multi-nucleation (Figure 3C and Supplemental Movies 4 and 5; for control see Figure 3B and Supplemental Movie 3). The shCLIC4 cells that did divide exhibited an extreme “blebbing” phenotype during anaphase. (Figure 3C, D arrows, F). The formation of blebs during telophase has been previously observed and has been suggested to function as a pressure release mechanism at the end of furrowing ^26–29^. These blebs typically occur at the poles of the dividing cell and are rarely observed at the furrow. In shCLIC4 cells, enlarged blebs could be observed at the furrow as well as poles of the cell, especially during the transition from the contracting cytokinetic ring to stable intercellular bridge (Figure 3C, D, F and Supplemental Movies 4 and 5). Since local changes in the cortical actin network are responsible for furrow ingression, cortical rigidity, and bleb formation and pressure release during anaphase, we imaged the anaphase cortical actin network using SiR-actin in live cells. We found that shCLIC4 cells had decreased levels of SiR-actin in both the furrow and poles of the cells, lending to the idea that these cells have a weaker cortical network and are therefore more likely to have an extreme blebbing phenotype (Figure 3G).

To further test whether CLIC4 is required for the cell division we used a combination of time-lapse microscopy and an optogenetic trap (termed ‘Mitotrap’) assay. In this assay, HeLa cells expressing endogenously tagged GFP-CLIC4 were transfected with Cry2 GFP-VHH and Tom20-CIB-stop (Figure 4A). Cells that receive both constructs will recruit endogenous GFP-CLIC4 to mitochondria when pulsed with a 488 nm laser (Figure 4B), thereby depleting CLIC4 from its normal localization at the plasma membrane. As shown in Figure 4C-E, cells that have CLIC4 anchored to the mitochondria also exhibit blebbing phenotypes, increase in mitosis time and, in some cases, regression of the cytokinetic furrow at late anaphase (see also Supplemental Movies 6 and 7). Taken together, these data indicate that CLIC4 is an important player in the fidelity of cell division, and likely plays a role in formation of the stable intercellular bridge and in regulating cytoskeletal remodeling.

**Figure 4.**
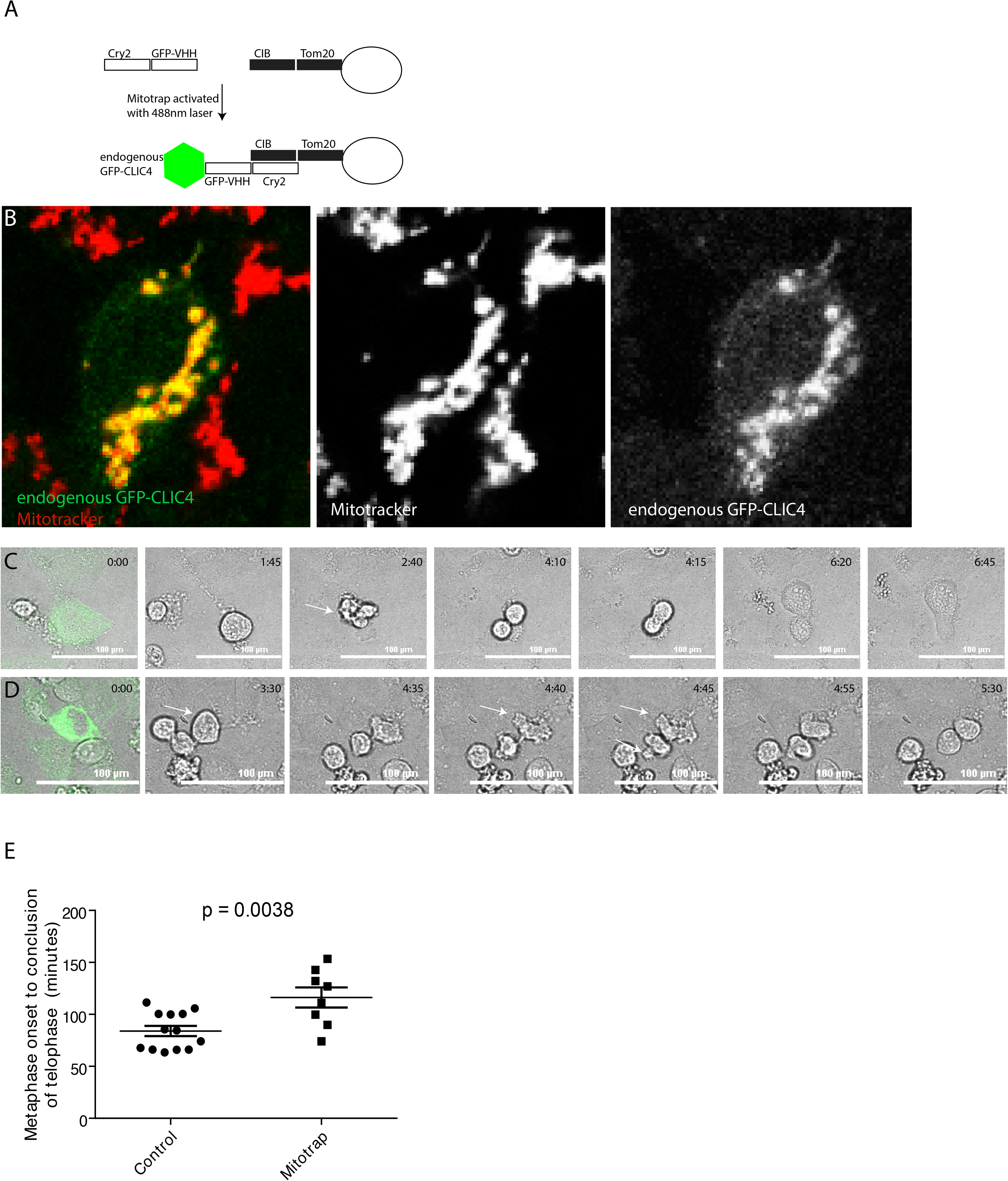
An optogenetic targeting of CLIC4 to the mitochondria delays cell division. (A) Schematic of Mitotrap used in this experiment. In all cases cells expressing endogenously tagged GFP-CLIC4 were co-transfected with Cry2-GFP-VHH and CIB-Tom20 plasmids and pulsed with a 488nm laser to re-target GFP-CLIC4 at the mitochondria. (B) Interphase cell exposed to 488nm to activate Mitotrap. Mitochondria is labeled in red and endogenous GFP-CLIC4 in green. (C-D) Still images from time-lapse microscopy where Mitotrap was activated. Arrows point to blebs or cytokinesis failure induced by GFP-CLIC4 Mitotrap. (E) Quantification of time required for cells to complete mitotic cell division in control and Mitotrapped cells. Data shown are the means and standard deviations.

### CLIC4 is required for the recruitment of non-muscle myosin IIA and IIB to the cytokinetic furrow

Our data demonstrate that CLIC4 is localized to the cytokinetic furrow and the absence of CLIC4 leads to increased blebbing and furrow regression, presumably due to the failure to transition from contractile actomyosin ring to ICB. Next, we decided to identify the mechanisms that mediate CLIC4 function during cell division. Recent studies identified several proteins involved in the regulation of mitotic blebbing process, namely the non-muscle myosins IIA (NMYIIA) and IIB (NMYIIB) ^30–32^. Furthermore, NMYIIA and NMYIIB have previously been shown to be responsible for regulating cortical rigidity during anaphase and telophase ^33^. We first tested whether CLIC4 contributes to recruitment and/or maintenance of NMYIIA and NMYIIB at the cleavage furrow. Consistent with this idea, GFP-CLIC4 colocalizes with NMYIIB during anaphase (Figure 5A). We measured NMYIIA and NMYIIB levels in the furrow in control and shCLIC4 cells and found that CLIC4 depletion leads to a decrease in NMYIIA and NMYIIB levels at the furrow during cytokinesis (Figure 5B-G), suggesting that CLIC4 may contribute to the spatiotemporal control of NMYIIA and NMYIIB during telophase.

**Figure 5.**
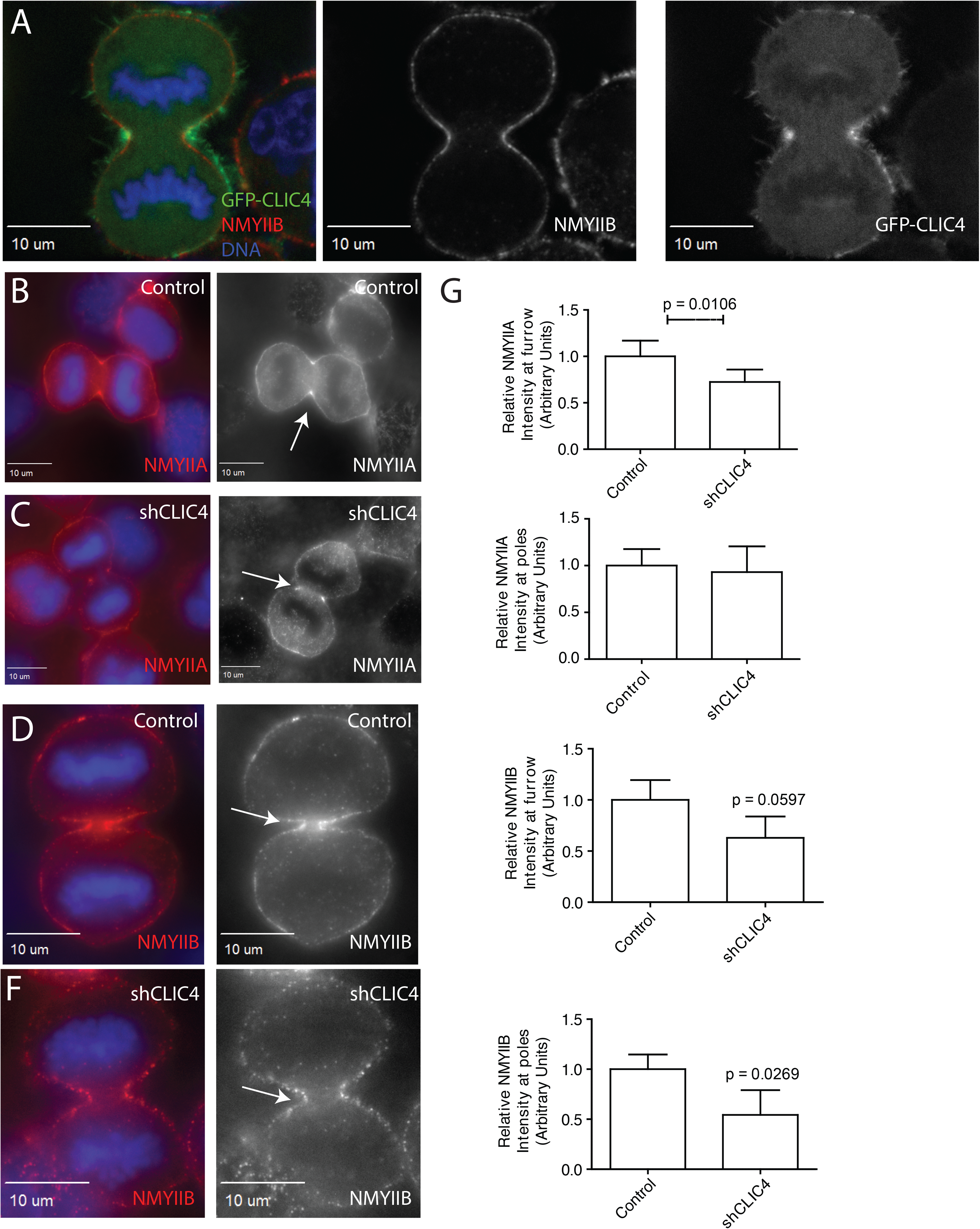
Loss of CLIC4 results in decreases in NMYIIA/B at cortex and furrow. (A) HeLa cells expressing GFP-CLIC4 were fixed and stained with anti-NMYIIB antibodies. (B-F) Control (B and D) or shCLIC4 (C and F) HeLa cells were fixed and-stained with either anti-NMYIIA (B and C) or anti-NMYIIB (D and F) antibodies. Arrows point to ingressing furrow during anaphase. (G) Quantification of NMYIIA or NMYIIB at the furrow or at the poles of the anaphase cell. The data shown are the means and standard deviations of 20-40 randomly picked cells from three different experiments.

### CLIC4 regulates phospho-ezrin recruitment to the plasma membrane

An additional mediator of cortical rigidity during cell division is ezrin, a member of the ERM family of proteins. There have been previous reports of CLIC4 and ezrin functional interactions. For example, it was shown that loss of CLIC4 in kidney and glomerular endothelium modulate activated ezrin levels ^16^. Furthermore, it has been shown that loss of moesin in Drosophila led to mitotic defects and a propensity for cells to bleb during anaphase ^2,3^. To begin examining a potential CLIC4 and ezrin functional interaction, we first analyzed phospho-ezrin localization during cell division and found that GFP-CLIC4 and phospho-ezrin colocalize during telophase (Figure 6A, see arrow). Next, we examined phospho-ezrin levels in shCLIC4 cells. We found that CLIC4 depletion led to a decrease in phospho-ezrin levels at the cell cortex and ingression furrow, suggesting that CLIC4 is necessary for the phosphorylation and activation of ezrin at the cytokinetic furrow (Figure 6 B-D). To further confirm that CLIC4 is required for proper phospho-ezrin localization at the cell cortex, we used our Mitotrap approach to acutely inactivate all cellular CLIC4 (Figure 6E-F). To that end, cells expressing endogenous GFP-CLIC4 were transfected with the Mitotrap constructs and exposed to 488 nm laser to “trap” CLIC4 at the mitochondria. Cells were then fixed and stained for phospho-ezrin. As shown in Figure 6G-H, inactivation of CLIC4 led to decrease in the amount of phospho-ezrin at the plasma membrane.

**Figure 6.**
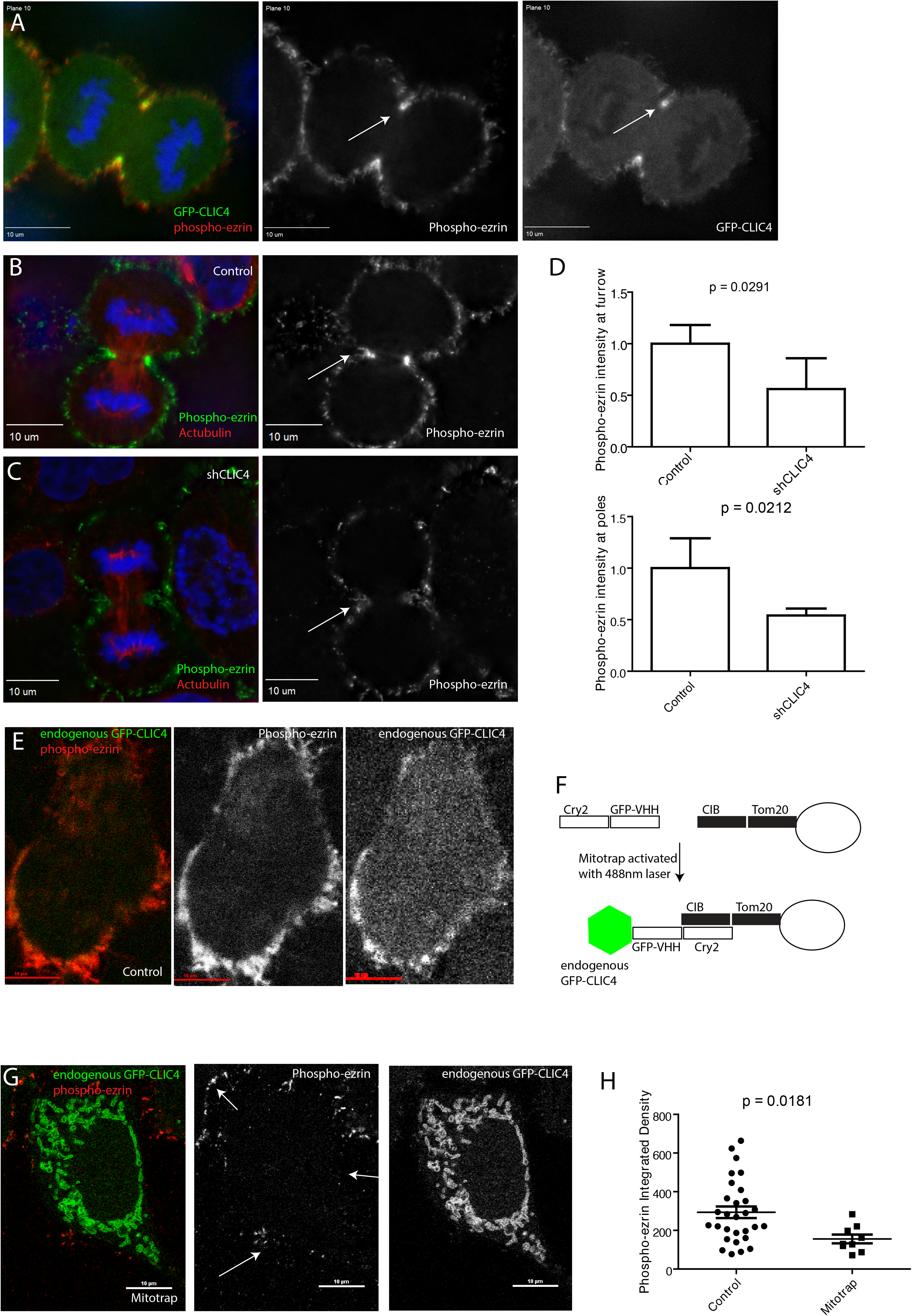
CLIC4 is necessary for efficient phospho-ezrin recruitment to the cleavage furrow. (A) Hela cells expressing GFP-CLIC4 were fixed and stained with anti-phospho-ezrin antibodies. Arrows point to the cleavage furrow. (B-D) Control or shCLIC4 cells were fixed and stained with anti-acetylated tubulin and anti-phospho-ezrin antibodies. Arrows point to the furrow. Panel (D) shows quantification of phospho-ezrin signal intensity in cleavage furrow and poles of the cell. Data shown are the means and standard deviations derived from three different experiments. (E-G) The GFP-CLIC4 Mitotrap assay designed to test the effect of CLIC4 depletion on the levels of Phospho-ezrin at plasma membrane. Panel (E) shows schematic of Mitotrap set up, where endogenous GFP-CLIC4 cells are transiently transfected with the Mitotrap plasmids and pulsed with a 488nm laser to trap CLIC4 at the mitochondria. Panels (E and G) shows localization of endogenously tagged GFP-CLIC4 and phospho-ezrin in cells before and after exposure to 488 nm wavelength pulse. (H) Quantification of phospho-ezrin levels at the plasma membrane before and after exposure to 488 nm to activate GFP-CLIC4 Mitotrap. Data shown are the means and standard deviations derived from three different experiments.

To further confirm that CLIC4 is involved in regulating phospho-ezrin targeting to the cytokinetic furrow, we next knocked-out CLIC4 in MDCK cells. MDCK cells are a canine kidney epithelial cell line that retained the ability to form polarized epithelial monolayers (Supplemental Figure 4A). Consistent with our data using HeLa cells, GFP-CLIC4 also localizes to the cleavage furrow in MDCK polarized monolayers (Supplemental Figure 4B). Unlike HeLa cells, dividing MDCK cells do not form blebs, likely due to the fact that they are still part of the epithelial monolayer and maintain tight junction and adherens junctions with neighboring interphase cells (Supplemental Figure 4B). Importantly, MDCK CLIC4-KO cells in telophase formed multiple membranous protrusions (Supplemental Figure 4C), suggesting CLIC4 is also required to maintain the rigidity of sub-plasma membrane cytoskeleton in epithelial cells. Finally, CLIC4-KO also led to a dramatic decrease in phospho-ezrin levels at the cytokinetic furrow in MDCK cells (Supplemental Figure 4D).

### MST4 kinase is a CLIC4-binding protein could regulate ezrin activation during anaphase

Since our data demonstrate the requirement of CLIC4 for successful completion of cytokinesis, we next decided to identify the molecular machinery governing CLIC4 function during mitosis. To that end, we used HeLa cells stably expressing pLVX:GFP-CLIC4 and performed immuno-precipitation using a GST-tagged anti-GFP nanobody followed by mass spectrometry in order to identify potential binding partners of CLIC4 (Supplemental Table 1). Our proteomic analysis identified numerous actin regulators as putative CLIC4-interacting proteins (Figure 7A). Importantly, a putative CLIC4-interacting protein identified in our proteomics was MST4, one of the kinases responsible for phosphorylating and activating ezrin, affecting the formation/activation of cortical actomyosin network ^34^. The *Dictyostelium discoideum* homolog of MST4 was shown to be necessary for cell division ^35^. Furthermore, recent work in *C. elegans* has shown that MST4 (*C. elegans* GCK-1) localizes to the contractile ring ^36^, and that depletion of MST4/GCK1 leads to collapse of intercellular canals in these animals. These data raise the possibility that MST4 may be targeted and/or activated by CLIC4 at the cytokinetic ring. To test this hypothesis, we first used glutathione bead pull-down assay to demonstrate that recombinant purified GST-CLIC4 binds to MST4 in cell lysates (Figure 7B). Since MST4 localization during mammalian cell division was never investigated, we next examined the localization of MST4 in mammalian cells throughout mitosis and found it to be enriched at the cytokinetic contractile ring (Figure 7C, see arrow). Interestingly, MST4 translocates to the furrow only at late telophase. This coincides with the timing of the formation of blebs outside the furrow, the process that appears to be regulated by CLIC4. To test whether MST4 recruitment to the furrow is mediated by CLIC4 we analyzed MST4 localization in shCLIC4 cells. Depletion of CLIC4 results in a moderate but significant decrease in MST4 at the furrow in late telophase (Figure 7D-E).

**Figure 7.**
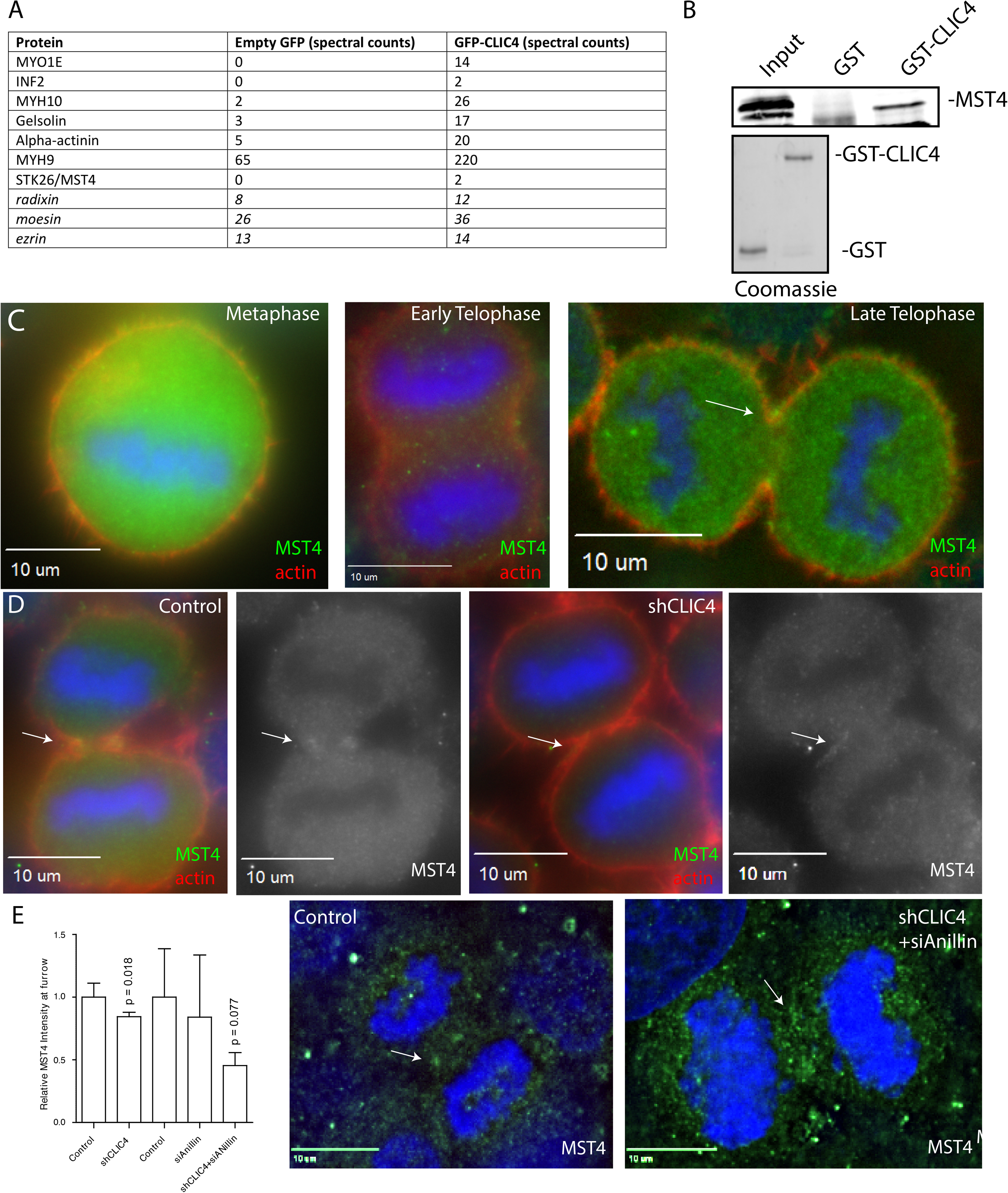
CLIC4 interacts with MST4 and affects its localization during anaphase. (A) Putative candidates revealed via immunoprecipitation of GFP-CLIC4 followed by mass spectrometry. (B) Glutathione bead pulldown assay using HeLa cell lysate and either GST only or GST-CLIC4. Coomassie staining showing equal loading as well as quality of the recombinant proteins used in pulldown assay. (C) HeLa cells were fixed and stained with phalloidin-Alexa568 and anti-MST4 antibodies. Arrow points to MST4 at the cleavage furrow. (D) Control or shCLIC4 HeLa cells were fixed and stained with anti-MST4 antibody. Arrows point to cleavage furrow. (E) Quantification of MST4 intensities in the furrow (top) and poles (bottom) in control, shCLIC4, siAnillin and shCLIC4/siAnillin cells. Data shown are the means and standard deviations of three individual experiments.

While depletion of CLIC4 did decrease the amount of MST4 at the furrow during late anaphase, the effect is quite moderate. We wondered whether other furrow proteins may also contribute to MST4 targeting during cell division. Recent work in *C. elegans* embryos suggested that anillin may bind to MST4 and mediate MST4 recruitment to cytokinetic furrow ^36^. To test whether anillin has a similar function in mammalian cells, we analyzed the localization of MST4 during anaphase in HeLa cells transfected with anillin siRNA. As shown in Figure 7E, anillin depletion alone had little effect on MST4 recruitment during cell division. Intriguingly, co-knock down of both anillin and CLIC4 resulted in a much greater decrease in MST4 localization at the cytokinetic furrow than knock-down of either of these proteins alone (Figure 7E). These findings suggest that in mammalian cells CLIC4 likely plays a primary role in mediating MST4 targeting during late anaphase. However, it is clear that anillin is also involved in MST4 targeting, especially in the context of CLIC4 depletion. Taken together, these data suggest that depletion of CLIC4 specifically at the cell cortex also results in loss of activated ezrin, providing a potential explanation for the increase in blebbing and failure in cell division.

## Discussion

Cytokinesis is an incredibly complex process that must be regulated appropriately in order to maintain mitotic fidelity. The actin cytoskeleton plays a key role during cytokinesis, forming the cytokinetic contractile ring that drives the separation of the mother cell into two daughter cells. Additionally, it is becoming well-established that the cortical actomyosin network is very important for successful completion of cytokinesis. Dynamic changes in the cortical actin cytoskeleton allows dividing cells to undergo dramatic shape changes, from rounding up during metaphase to the elongation during anaphase and finally flattening during late telophase and abscission (Figure 8). Additionally, at the end of cleavage furrow ingression, the cytokinetic contractile ring needs to be remodeled and converted to an actin network that stabilizes the newly formed intercellular bridge until the abscission step of cytokinesis. The failure of intercellular bridge stabilization often leads to regression of cleavage furrow, leading to polyploidy. Interestingly, in some developmental instances, the abscission step is delayed, leading to the formation of stable syncytial connections between cells, also known as ring canals ^37–40^. How the cortical actin cytoskeleton coordinates with the cytokinetic contractile ring to simultaneously allow various cell shape changes as well as furrow ingression is still under investigation by our lab and others. In this study, we have identified CLIC4 as a new component of cytokinetic furrow that plays a key role in regulating the actomyosin cytoskeleton and maintaining this network during cell division.

**Figure 8.**
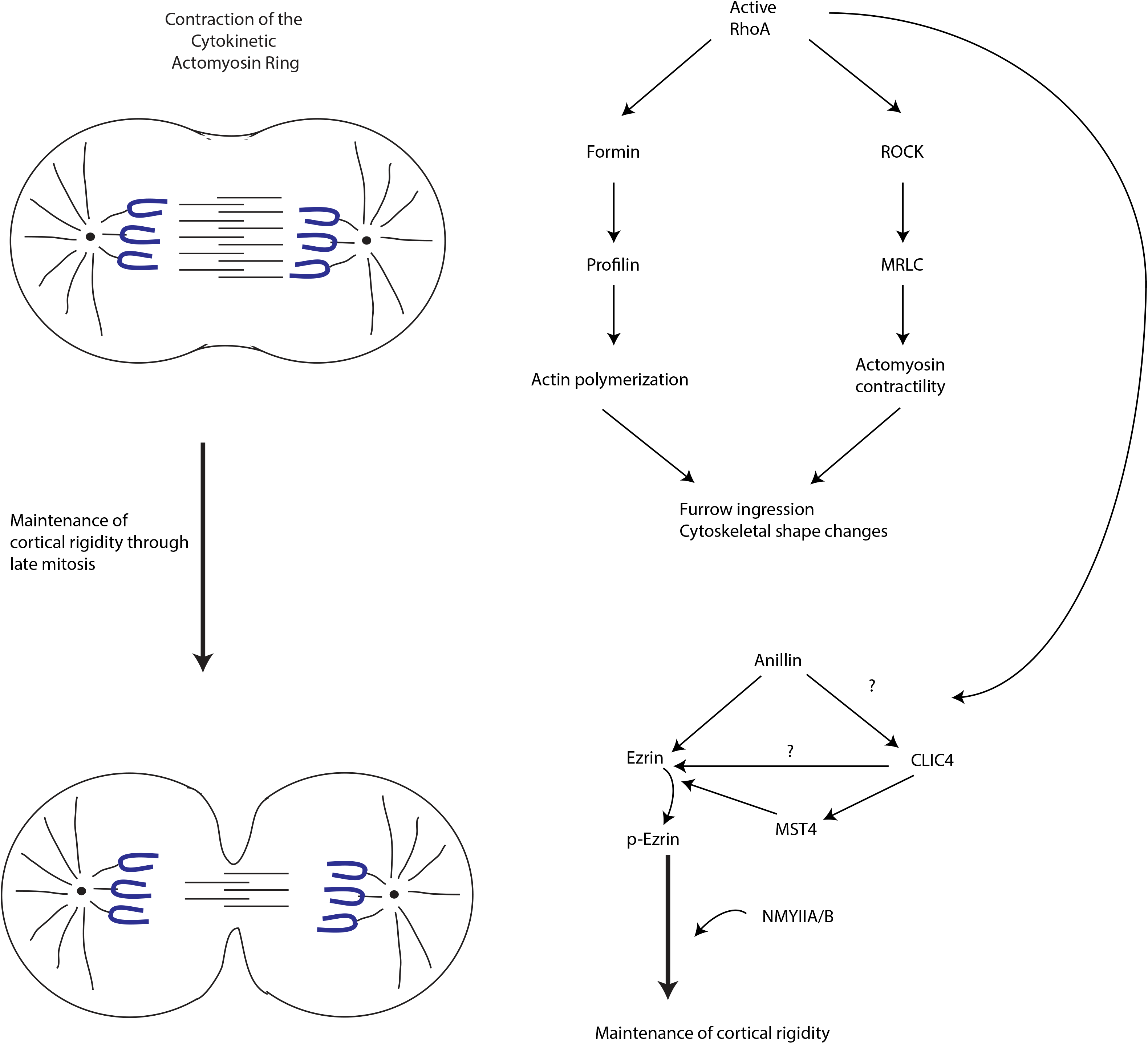
Model for CLIC4 during the anaphase-to-telophase transition.

We originally identified CLIC4 as putative midbody-associated protein from our midbody proteomics analysis^18^ and have confirmed that CLIC4 is indeed present at the midbody during late telophase, consistent with a previous study ^21^. Thus, we initially studied CLIC4 as a putative regulator of abscission. However, we found no evidence that CLIC4 is directly required for abscission (data not shown). In contrast, our localization analysis suggested that CLIC4 is actually removed from the abscission site (Figure 1B), although the mechanism mediating CLIC4 exclusion during abscission remains unclear. Since it has been shown that actin filaments need to be removed from the abscission site^22,23^ and CLIC4 is known to regulate actin polymerization, it is tempting to speculate that CLIC4 exclusion from the abscission site may contribute to the regulation of determining the timing and location of the abscission.

Our previously published midbody proteome identified a number of proteins that may be involved in regulating actin dynamics during furrow ingression. Since CLIC4 has been implicated in regulating actin dynamics in interphase cells, we have explored the possibility that CLIC4 may regulate actin dynamics during cytokinesis and have identified CLIC4 as a new component of cytokinetic furrow (Figure 1). We show that CLIC4 localizes to the cleavage furrow during the onset of cytokinetic ring contraction and that CLIC4 recruitment is dependent on RhoA activation (Figure 2). We propose that in addition to activating NMYlIA/IIB and inducing formin-dependent actin polymerization, RhoA also drives the accumulation of CLIC4 at the furrow (Figure 8).

Following the discovery that CLIC4 is a component of the cytokinetic furrow, we next asked what role CLIC4 plays during cytokinesis. Upon depletion of CLIC4, we found an increase in the percentage of multinucleated cells as well as an increase in division times (Figure 3), phenotypes typically observed as a result of failure in cytokinetic ring contraction. Surprisingly, our time-lapse analysis demonstrated that CLIC4 depletion has little effect on the furrow formation and ingression. Instead, some shCLIC4 cells failed to convert the ingressing furrow to a stable intercellular bridge. Furthermore, cells that managed to divide exhibited a high blebbing phenotype (Figure 3C, D, F). Blebs are commonly observed during late anaphase and are believed to serve the role of pressure release valves due to rapid decrease in cell volume ^26,27,33,41^. However, the burst of blebbing is usually brief and restricted to the poles of the dividing cell. In contrast, shCLIC4 cells form much larger blebs, with many of them occurring at the furrow (Figure 3C, D, F), suggesting that CLIC4 may predominately function to regulate the cortex stiffness and inhibit blebbing at the furrow during its conversion to the stable intercellular bridge. Consistent with this, we observed decreased intensity of the cortical actin network in shCLIC4 cells in telophase (Figure 3G). Since NMYIIA/IIB were previously shown to regulate bleb formation ^31–33,41^, we also examined the recruitment of NMYIIA /IIB in shCLIC4 cells. Consistent with the involvement of CLIC4 in the regulation of blebbing, loss of CLIC4 resulted in the decrease of the NMYIIA/IIB at the furrow cortex in late anaphase (Figure 5). Based on these data we propose that CLIC4 regulates furrow cortex stiffness, and consequently blebbing, during anaphase. We also speculate that CLIC4-dependent machinery plays a key role during the formation and stabilization of intercellular bridge (Figure 8).

CLIC4 has been initially described as a putative chloride channel, but recent works have determined it has other functions, including the regulation of cytoskeletal dynamics. Indeed, CLIC4 has been thought to regulate actin polymerization in several different contexts and appears to predominately enhance formation of formin-induced actin polymerization while inhibiting Arp2/3-dependent formation of branched actin networks ^14,15^. What remains unclear is how CLIC4 regulates actin dynamics since it does not affect actin polymerization directly, nor does it bind to directly to polymerized actin filaments ^13,42^. Some recent work suggested that CLIC4 may bind to profilin^14^ although it remains unclear whether this interaction plays any role in regulating actin during cell division. In this work we identified MST4 as a CLIC4-interacting protein and have shown that CLIC4 contributes to mediating the recruitment of MST4 to the cytokinetic contractile ring (Figure 7, 8). Interestingly, while CLIC4 is enriched at the cytokinetic contractile ring from the onset of anaphase, MST4 is only recruited there at late anaphase. This is consistent with the putative role of MST4 in regulating actin cytoskeleton remodeling during the formation of stable intercellular bridge. These studies in *C. elegans* have pointed to a role for MST4/GCK-1 in stabilizing intercellular ring canals during development, and MST4/GCK-1 was shown to be localized to the contractile ring, confirming our mammalian localization ^36,39^.

MST4 is one of the kinases capable of phosphorylating ezrin. The role of activated ezrin in mediating the attachment of actin filament to the plasma membrane is now well-defined^43^ and seems to depend on the ability of ezrin to co-bind filamentous actin and phosphatidyl-inositol 4,5 di-phosphate (PIP2), as well as several plasma membrane proteins. Importantly, PIP2 is required for successful completion of cytokinesis and was shown to be enriched at the intracellular bridge ^44^. It is possible that the recruitment of MST4 to the furrow at late anaphase ensures the stabilization of filamentous actin and its attachment to the plasma membrane in the forming intracellular bridge (Figure 8). Similarly, activation of this CLIC4-MST4-ezrin pathway would increase furrow stiffness, inhibiting blebbing at the furrow membranes. However, we observed only a slight effect on MST4 recruitment during anaphase in shCLIC4 cells. Yet, we saw a much larger effect on phospho-ezrin levels in the furrow in HeLa-shCLIC4 and MDCK-CLIC4-KO cells. This suggests that an alternative mechanism for CLIC4 in modulating phospho-ezrin expression/localization during anaphase. One obvious mechanism would be for a direct CLIC4-ezrin binding interaction. However, ezrin did not pass our cutoff in our mass spectrometry data and we could not demonstrate that CLIC4 binds to ezrin (Supplemental Figure 3). While we cannot fully rule out that CLIC4 binds directly to ezrin, it is likely that in addition to MST4, CLIC4 may also recruit or regulate some other phospho-ezrin regulator.

We also have shown that CLIC4 contributes to NMYIIA/IIB recruitment to the furrow. However, how CLIC4 regulates NMYIIA/IIB remains unclear. We did not pull-down any kinases that might be responsible for phosphorylation and activation of NMYIIA/IIB, but we cannot rule out the possibility that CLIC4 can regulate myosin in this fashion. Alternatively, CLIC4 may regulate NMYIIA/IIB indirectly by modulating cortical actin cytoskeleton.

We conclude that CLIC4 is a novel component of the cytokinetic furrow that regulates cortical rigidity during cell division. We propose that CLIC4 functions, at least in part, by recruiting MST4 and regulating ezrin activation at the furrow plasma membrane (Figure 8). However, many questions remain. Does CLIC4 regulate recruitment of other actin regulators? What are the roles, if any, other CLIC family members during cell division? Does CLIC4 have similar functions *in vivo*, especially in the context of formation of long-lived intracellular bridges or ring canals? Finally, what are the molecular mechanisms driving CLIC4-mediated recruitment of phospho-ezrin to the cell cortex? With the identification of a RhoA/CLIC4/MST4/Ezrin pathway, we and others can start answering these questions and systematically analyze the role for CLIC4 during cytokinesis.

## Acknowledgements

We would like to thank Ching-Hwa Sung for providing pCAG:IRES-CLIC4. This work was partially funded by R01s DK064380 and GM122768 to RP.

---

**Supplemental Figure 1. GFP-CLIC4 localization during anaphase and endogenous CLIC4 upon anillin knockdown**

(A) Exogenous GFP-CLIC4 is localized to cell cortex and actin-structures (such as filopodia) during interphase.

(B) Creation of endogenous GFP-CLIC4 in HeLa cells. Note the lack of CLIC4 band at 28kDa. Both lanes are endogenous GFP-CLIC4 lysates, 30 and 60 μg. Blot shown is original and uncropped.

(C) Knockdown of anillin with siRNA reduces the amount of endogenous GFP-CLIC4 at the cleavage furrow. Error bars represent standard deviation from three biological replicates, with at least 8 cells per replicate. P-value determined by unpaired Student’s t-test.

(D) HeLa cells expressing endogenous GFP-CLIC4 were transfected with the Cry2PHR-mCH-RhoA plasmid, incubated for 24 hours, and activated via pulse with 488nm laser. Cells were fixed and imaged to demonstrate clustering and activation of RhoA is sufficient to relocalize CLIC4.

**Supplemental Figure 2. Validation of shCLIC4 and IAA94 treatments**

(A) Efficacy of CLIC4 shRNA. Left, western blot of shCLIC4 (left) and control (right) lysates. Right, quantification of three biological replicates.

(B) Frequency of multinucleation in DMSO or IAA94 treated cells. Cells were treated with IAA94 for 24 hours, then fixed and stained with phalloidin and Hoechst.

(C-D) Endogenous GFP-CLIC4 localizes normally upon treatment with IAA94. Cells were treated with DMSO or IAA94 for 24 hours, fixed, and stained with phalloidin 568. Intensity of endogenous GFP-CLIC4 was measured at the furrow.

**Supplemental Figure 3. GST-CLIC4 does not bind to ezrin.**

(A) HeLa cells were lysed and 1mg was subject to incubation with 10μg GST or GST-CLIC4. Glutathione beads were added, protein was eluted with 1x SDS, and eluate were subject to SDS-PAGE separation followed by western blot for ezrin. Blot shown is original, uncropped blot.

**Supplemental Figure 4. CLIC4 regulates phospho-ezrin recruitment to the cytokinetic furrow in MDCK cells**

(A) Western blot of cell lysates derived from control or CLIC4-KO MDCK cells.

(B) MDCK cells expressing GFP-CLIC4 were grown on collagen-coated filters to allow full polarization and then stained with phalloidin-Alexa568. Arrow points to cytokinetic cleavage furrow.

(C) MDCK or MDCK-CLIC4-KO cells were grown on collagen-coated filters. Cells were then fixed and stained with phalloidin and anti-acetylated tubulin (central spindle marker) antibodies. Arrows point to membranous protrusions in CLIC4-KO cells.

(D) MDCK or MDCK-CLIC4-KO cells were grown on collagen-coated filters. Cells were then fixed and stained with phalloidin and anti-phospho-ezrin antibodies. Arrows point to cytokinetic furrow.

**Supplemental Figure 5. Original, uncropped western blots.**

(A) Original western blot for HeLa shCLIC4 cell line

(B) Original western blot for MST4 pulldown.

(C) Original western blot for MDCK-KO cell line.

**Supplemental Movie 1.** 3D volume reconstruction of cells expressing exogenous GFP-CLIC4 (green) and stained with anti-acetylated tubulin (red).

**Supplemental Movie 2.** Time lapse microscopy of cells expressing exogenous GFP-CLIC4 during anaphase.

**Supplemental Movie 3.** Time lapse microscopy of representative control cells undergoing cell division.

**Supplemental Movie 4.** Time lapse microscopy of shCLIC4 cells undergoing cell division, exhibiting extreme blebbing phenotypes and delayed mitosis.

**Supplemental Movie 5.** Time lapse microscopy of shCLIC4 cell undergoing cell division, exhibiting extreme blebbing, delayed mitosis and failing to divide.

**Supplemental Movie 6.** Time lapse microscopy of Mitotrap cell undergoing cell division, exhibiting extreme blebbing and delayed mitosis.

**Supplemental Movie 7.** Time lapse microscopy of Mitotrap cell undergoing cell division, exhibiting a failure to divide.

**Supplemental Table 1.** Complete list of proteins pulled down in GFP-CLIC4 immunoprecipitation followed by mass spectrometry.

## References

1. Lancaster, O. M. & Baum, B. Shaping up to divide: Coordinating actin and microtubule cytoskeletal remodelling during mitosis. Semin. Cell Dev. Biol. 34, 109–115 (2014).

2. Kunda, P., Pelling, A. E. & Liu, T. Article Moesin Controls Cortical Rigidity, Cell Rounding, and Spindle Morphogenesis during Mitosis. 91–101 (2008). doi:10.1016/j.cub.2007.12.051

3. Carreno, S., Kouranti, I., Glusman, E. S., Fuller, M. T. & Echard, A. Moesin and its activating kinase Slik are required for cortical stability and microtubule organization in mitotic cells. 180, 739–746 (2008).

4. Mabuchi, I. & Okuno, M. THE EFFECT DIVISION OF MYOSIN ANTIBODY ON THE OF STARFISH Antiserum against starfish egg myosin was produced in rabbits. Antibody specific- ity to myosin was demonstrated by Ouchterlony - s immunodiffusion test and by immunoelectrophoresis in the presenc. J. Cell Biol. 74, 251–263 (1977).

5. Fujiwara, K. & Pollard, T. D. Fluorescent antibody localization of myosin in the cytoplasm, cleavage furrow, and mitotic spindle of human cells. J. Cell Biol. 71, 848–875 (1976).

6. Wagner, E. & Glotzer, M. Local RhoA activation induces cytokinetic furrows independent of spindle position and cell cycle stage. J. Cell Biol. 213, 641–649 (2016).

7. Piekny, A. J. & Glotzer, M. Anillin Is a Scaffold Protein That Links RhoA, Actin, and Myosin during Cytokinesis. Curr. Biol. 18, 30–36 (2008).

8. Straight, A. F., Field, C. M. & Mitchison, T. J. Anillin Binds Nonmuscle Myosin II and Regulates the Contractile Ring. Mol. Biol. Cell (2005). doi:10.1091/mbc.e04-08-0758

9. Sun, L. et al. Mechanistic Insights into the Anchorage of the Contractile Ring by Anillin and Mid1. Dev. Cell 33, 413–426 (2015).

10. Hiruma, S., Kamasaki, T., Otomo, K., Nemoto, T. & Uehara, R. Dynamics and function of ERM proteins during cytokinesis in human cells. 591, 3296–3309 (2017).

11. Roubinet, C., Decelle, B., Dorn, J. F. & Payrastre, B. Molecular networks linked by Moesin drive remodeling of the cell cortex during mitosis. 195, 99–112 (2011).

12. Hickson, G. R. X., Echard, A. & O’Farrell, P. H. Rho-kinase controls cell shape changes during cytokinesis. Curr. Biol. 16, 359–370 (2006).

13. Argenzio, E. et al. CLIC4 regulates cell adhesion and 1 integrin trafficking. J. Cell Sci. 127, 5189–5203 (2014).

14. Argenzio, E. et al. Profilin binding couples chloride intracellular channel protein CLIC4 to RhoA-mDia2 signaling and filopodium formation. J. Biol. Chem. 293, 19161–19176 (2018).

15. Chou, S. Y. et al. CLIC4 regulates apical exocytosis and renal tube luminogenesis through retromer- and actin-mediated endocytic trafficking. Nat. Commun. (2016). doi:10.1038/ncomms10412

16. Tavasoli, M. et al. Both CLIC4 and CLIC5A activate ERM proteins in glomerular endothelium. 945–957 (2018). doi:10.1152/ajprenal.00353.2016

17. Mangan, A. J. et al. Cingulin and actin mediate midbody-dependent apical lumen formation during polarization of epithelial cells. Nat. Commun. 7, 12426 (2016).

18. Peterman, E. et al. The post-abscission midbody is an intracellular signaling organelle that regulates cell proliferation. Nat. Commun. 10, 3181 (2019 ).

19. Skop, A. R., Liu, H., Yates, J., Meyer, B. J. & Heald, R. Dissection of the mammalian midbody proteome reveals conserved cytokinesis mechanisms. Science (80-.). 305, 61–66 (2004).

20. Capalbo, L. et al. The midbody interactome reveals unexpected roles for PP1 phosphatases in cytokinesis. Nat. Commun. (2019). doi:10.1038/s41467-019-12507-9

21. Berryman, M. A. & Goldenring, J. R. CLIC4 Is Enriched at Cell-Cell Junctions and Colocalizes With AKAP350 at the Centrosome and Midbody of Cultured Mammalian Cells. Cell Motil. Cytoskeleton (2003). doi:10.1002/cm.10141

22. Schiel, J. A. et al. FIP3-endosome-dependent formation of the secondary ingression mediates ESCRT-III recruitment during cytokinesis. Nat. Cell Biol. 14, 1068–78 (2012).

23. Dambournet, D. et al. Rab35 GTPase and OCRL phosphatase remodel lipids and F-actin for successful cytokinesis. Nat. Cell Biol. 13, 981–988 (2011).

24. Bugaj, L. J., Choksi, A. T., Mesuda, C. K., Kane, R. S. & Schaffer, D. V. Optogenetic protein clustering and signaling activation in mammalian cells. Nat. Methods (2013). doi:10.1038/nmeth.2360

25. Pertz, O., Hodgson, L., Klemke, R. L. & Hahn, K. M. Spatiotemporal dynamics of RhoA activity in migrating cells. Nature (2006). doi:10.1038/nature04665

26. Cattin, C. J. et al. Mechanical control of mitotic progression in single animal cells. Proc. Natl. Acad. Sci. 112, 11258–11263 (2015).

27. G.T. Charras. A short history of blebbing. J. Microsc. 231, 466–478 (2008).

28. Paluch, E., Piel, M., Prost, J., Bornens, M. & Sykes, C. Cortical actomyosin breakage triggers shape oscillations in cells and cell fragments. Biophys. J. 89, 724–733 (2005).

29. Sedzinski, J. et al. Polar actomyosin contractility destabilizes the position of the cytokinetic furrow. Nature (2011). doi:10.1038/nature10286

30. Wang, Kangji, Carsten Wloka, E. B. Non-muscle Myosin-II Is Required for the Generation of a Constriction Site for Subsequent Abscission Non-muscle Myosin-II Is Required for the Generation of a Constriction Site for Subsequent Abscission. ISCIENCE 13, 69–81 (2019).

31. Taneja, N. & Burnette, D. T. Myosin IIA drives membrane bleb retraction.

32. Taneja, N. et al. Precise Tuning of Cortical Contractility Regulates Mechanical Equilibrium During Cell Division. SSRN Electron. J. (2018). doi:10.2139/ssrn.3155577

33. Mitchison, T. J., Charras, G. T. & Mahadevan, L. Implications of a poroelastic cytoplasm for the dynamics of animal cell shape. Seminars in Cell and Developmental Biology (2008). doi:10.1016/j.semcdb.2008.01.008

34. ten Klooster, J. P. et al. Mst4 and Ezrin Induce Brush Borders Downstream of the Lkb1/Strad/Mo25 Polarization Complex. Dev. Cell 16, 551–562 (2009).

35. Rohlfs, M., Arasada, R., Batsios, P., Janzen, J. & Schleicher, M. The Ste20-like kinase SvkA of Dictyostelium discoideum is essential for late stages of cytokinesis. J. Cell Sci. 120, 4345–4354 (2007).

36. Rehain-Bell, Kathryn, Werner, Michael, Doshi, Anusha, Cortes, Daniel, Sattler, Adam, Vuong-Brender, Thanh, Labouesse, Michel, and Maddox, A. Novel cytokinetic ring components limit RhoA activity and contractility. Biorxiv (2019).

37. Haglund, K., Nezis, I. P. & Stenmark, H. Structure and functions of stable intercellular bridges formed by incomplete cytokinesis during development. Commun. Integr. Biol. 4, 1–9 (2011).

38. Haglund, K. et al. Cindr Interacts with Anillin to Control Cytokinesis in Drosophila melanogaster. Curr. Biol. 20, 944–950 (2010).

39. Rehain-Bell, K. et al. A Sterile 20 Family Kinase and Its Co-factor CCM-3 Regulate Contractile Ring Proteins on Germline Intercellular Bridges. Curr. Biol. 27, 860–867 (2017).

40. Amini, R. et al. C. Elegans Anillin proteins regulate intercellular bridge stability and germline syncytial organization. J. Cell Biol. 206, 129–143 (2014).

41. Charras, G. T., Hu, C., Coughlin, M. & Mitchison, T. J. Reassembly of contractile actin cortex in cell blebs. 175, 477–490 (2006).

42. Suginta, W., Karoulias, N., Aitken, A. & Ashley, R. H. Chloride intracellular channel protein CLIC4 (p64H1) binds directly to brain dynamin I in a complex containing actin, tubulin and 14-3-3 isoforms. Biochem. J. 359, 55–64 (2001).

43. Fehon, R. G., McClatchey, A. I. & Bretscher, A. Organizing the cell cortex: The role of ERM proteins. Nature Reviews Molecular Cell Biology (2010). doi:10.1038/nrm2866

44. Kouranti, I., Sachse, M., Arouche, N., Goud, B. & Echard, A. Rab35 Regulates an Endocytic Recycling Pathway Essential for the Terminal Steps of Cytokinesis. Curr. Biol. 16, 1719–1725 (2006).

