## Supplementary figures and images for "CLIC4 is a cytokinetic cleavage furrow protein that regulates cortical cytoskeleton stability during cell division"

### Supplemental Figures 1-4

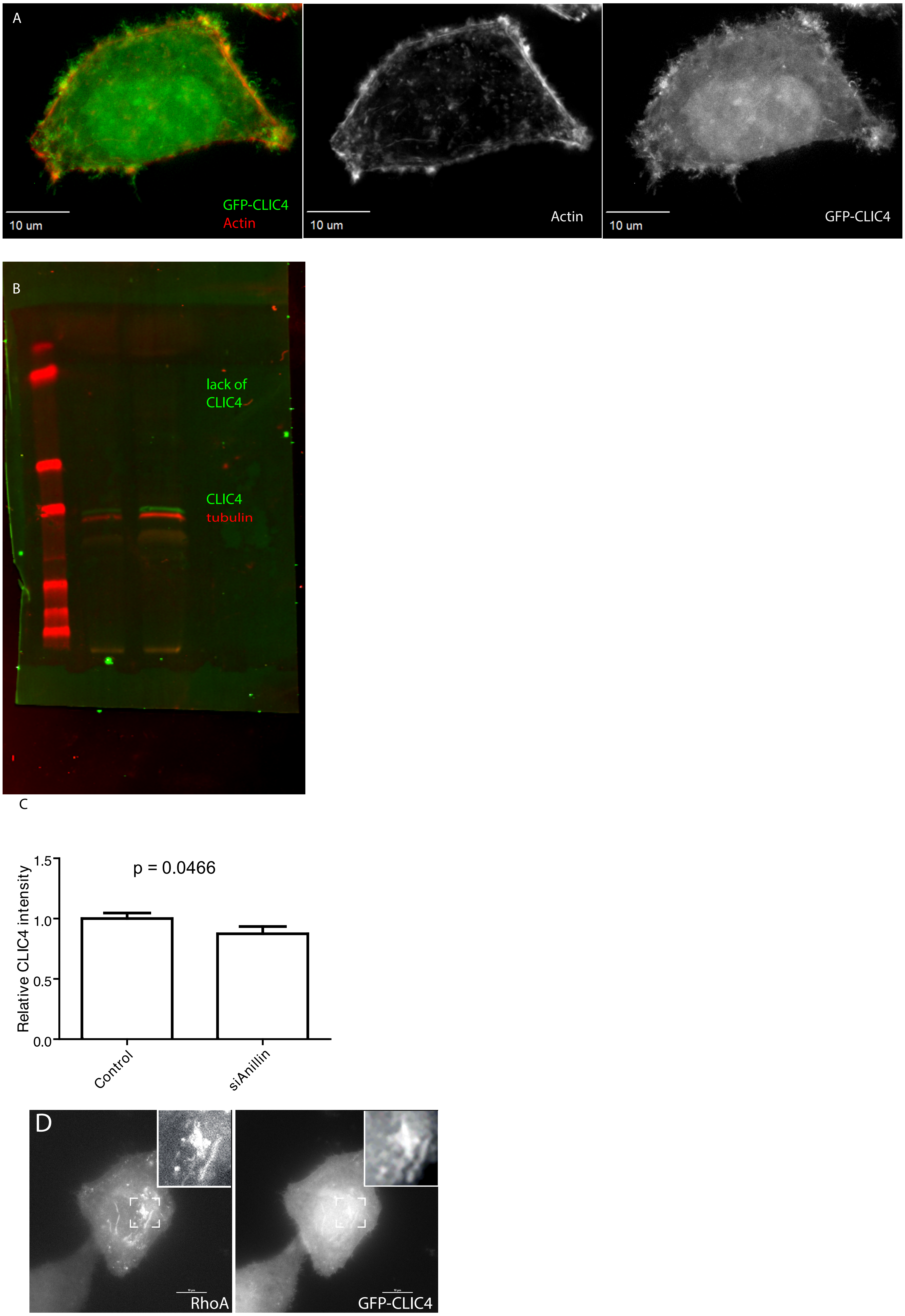

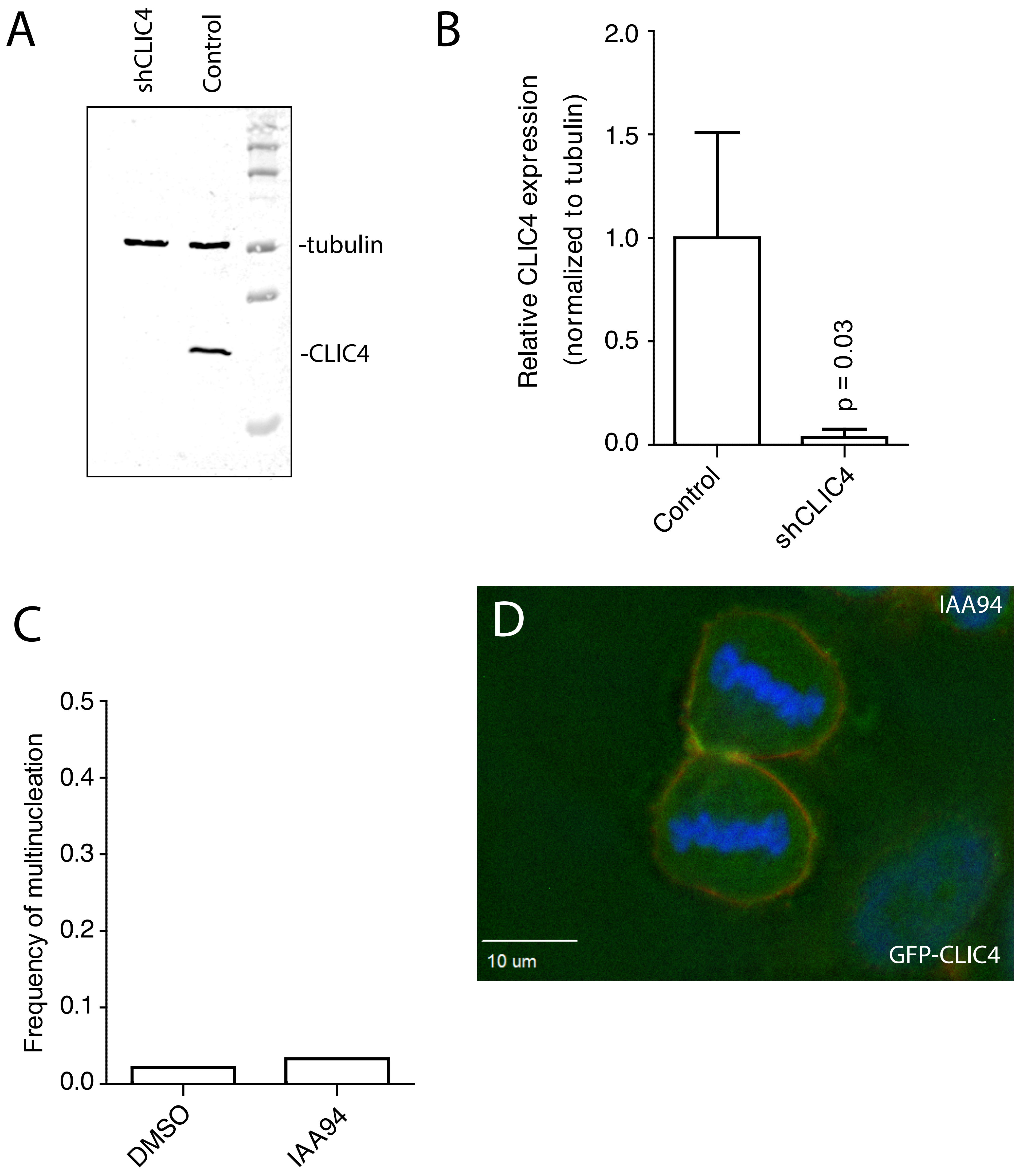

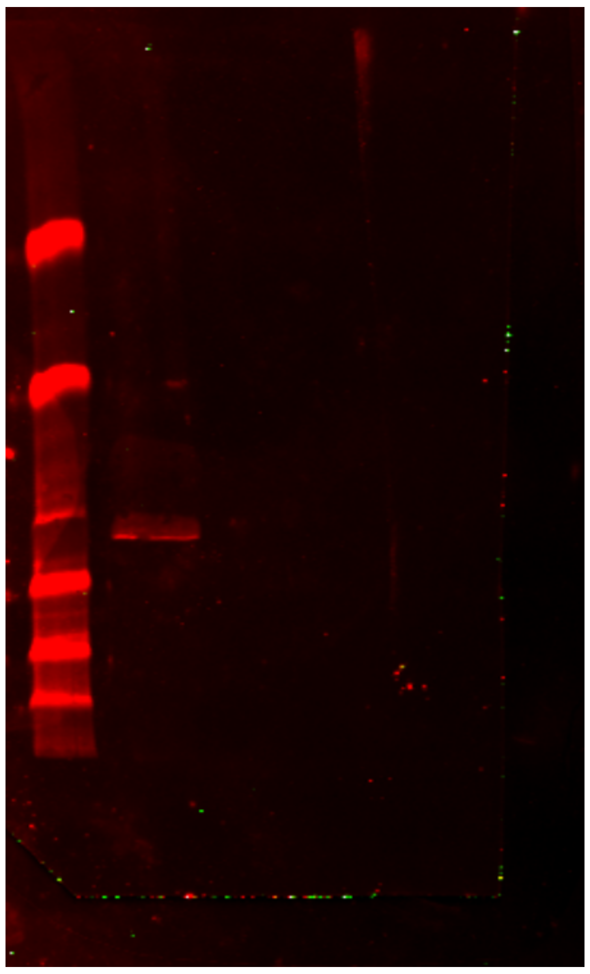

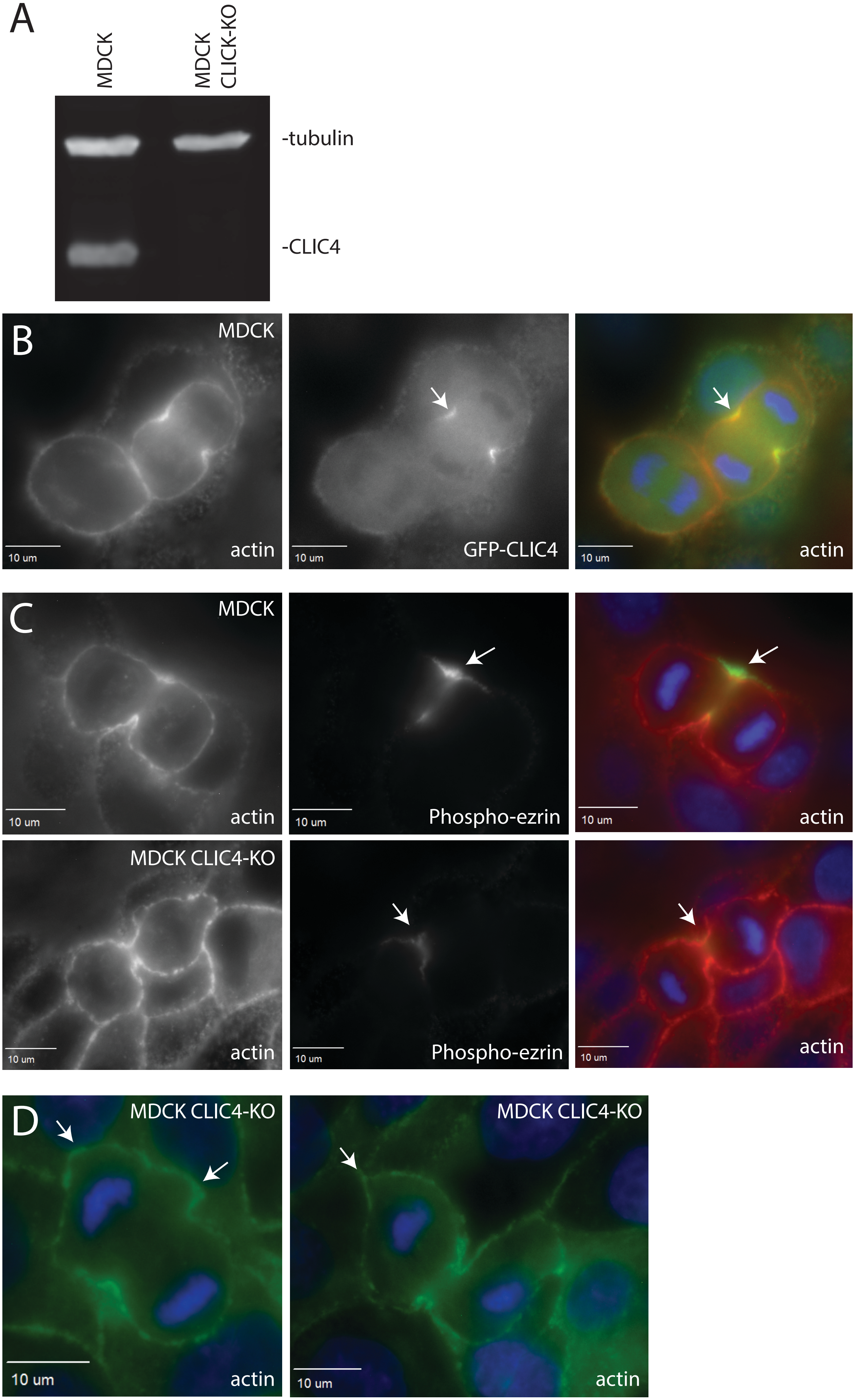
